# Genomic analyses identify biological processes in ZKSCAN3-deficient colorectal cancer cells

**DOI:** 10.1101/2021.12.30.474589

**Authors:** Zhiwen Qian, Tingxiang Chang, Tingting Zhang, Jing Wang, Hanming Gu

**Affiliations:** SHU-UTS SILC School, Shanghai University, Shanghai, China

## Abstract

Zinc finger with KRAB and SCAN domain 3 (ZKSCAN3) is associated with cell differentiation, cell proliferation and apoptosis, which has been reported as a critical driver of colorectal cancer. However, the mechanism and function of ZKSCAN3 in colorectal cancer is still unclear. Here, our objective is to identify the functional molecules and signaling by analyzing the RNA-seq data. The GSE172201 was created by the Illumina NovaSeq 6000 (Homo sapiens). The KEGG and GO analyses indicated the immune defense response to virus and transcription activity are major processes in the ZKSCAN3 KO colorectal cancer cells. Moreover, we determined ten key molecules including STAT1, MX1, DDX58, PPARG, EGFR, APP, BST2, DLG4, OASL, and IFIT2. Therefore, our study may provide the novel knowledge of ZKSCAN3 mediated colorectal cancer.

## Introduction

Colorectal cancer is one the most common cancers in the world, which has the highest incidence in Europe and North America^1^. The prognosis of colorectal cancer has increased during the past decades including Australia, US, Canada, and European countries^1–3^. There is no simple factor that causes colorectal cancer^4^. The known risk factors include age, sex, family history, obesity and diabetes^4^. The molecules underlying progression of this cancer are essential since they involve in the prognosis and treatment^5^. The relationship between the molecules and mechanism has become the key to treat the hereditary forms of colorectal cancer^6^.

ZKSCAN3 is a kind of oncogene that is increased in colon cancer frequently^7^. ZKSCAN3 contains the zinc finger structure and is a member of the family of KRAB and SCAN domain proteins^7^. This protein family plays a critical role in biological processes including apoptosis^8^. It was reported that ZKSCAN3 is an activator for the progression of colon cancer^9^.

In this study, we determined the biological functions of ZKSCAN3 in colorectal cancer by using the RNA-seq data. We identified a number of DEGs and biological processes and constructed the protein-protein interaction (PPI) network. The DEGs and PPI network may shed light on the progression mechanism of colorectal cancer.

## Methods

### Data resources

Gene dataset GSE172201 was downloaded from the GEO database. The data was produced by the Illumina NovaSeq 6000 (Homo sapiens) (Department of Pathology, Kyung Hee University, 26 Kyungheedaero, Seoul, South Korea). The analyzed dataset includes 3 groups of WT control HCT116 colorectal cancer cell line and 3 groups of ZKSCAN3 KO HCT116-1 colorectal cancer cell line.

### Data acquisition and processing

The data were organized and conducted by R package as previously described^10–14^. We used a classical t-test to identify DEGs with P< 0.05 and fold change ≥1.5 as being statistically significant.

The Kyoto Encyclopedia of Genes and Genomes (KEGG) and Gene Ontology (GO) KEGG and GO analyses were conducted by the R package (clusterProfiler) and Reactome. P<0.05 were considered statistically significant.

### Protein-protein interaction (PPI) networks

The Molecular Complex Detection (MCODE) was used to construct the PPI networks. The significant modules were produced from constructed PPI networks and String network. The pathway analyses were performed by using Reactome (https://reactome.org/), and P<0.05 was considered significance.

## Results

### Identification of DEGs of ZKSCAN3-deficient colorectal cancer cell line

To study the effects of ZKSCAN3 in colorectal cancers, we analyzed the RNA-seq data from the ZKSCAN3 knockout (KO) colorectal cancer cell line. A total of 139 genes were identified with the threshold of P < 0.05. The top up- and down-regulated genes in the ZKSCAN3 KO colorectal cancer cell line were identified by the heatmap and volcano plot (Figure 1). The top ten DEGs were listed in Table 1.

**Figure 1.**
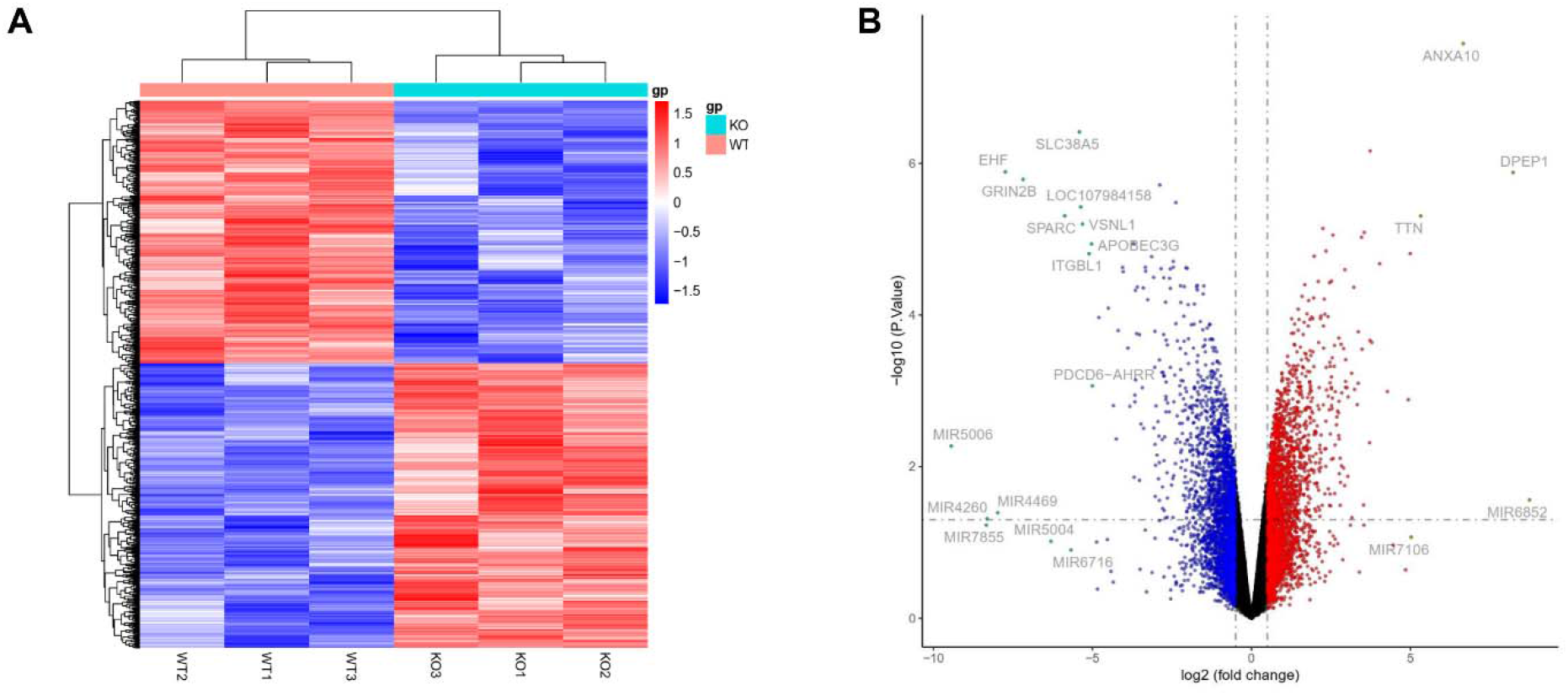
Heatmap and volcano plot were created from the WT and ZKSCAN3-deficient colorectal cancer cells. (A) Heatmap of significant DEGs. Significant DEGs (P < 0.01) were used to produce the heatmap. (B) Volcano plot for DEGs of the WT and ZKSCAN3-deficient colorectal cancer cells. The most significantly changed genes are highlighted by grey dots.

**Table 1.**

| **Entrez gene** | **Gene symbol** | **Fold-change** | **Regulation** |
| --- | --- | --- | --- |
| Top 10 down-regulated DEGs |  |  |  |
| 26298 | EHF | -7.739803829 | Down |
| 2904 | GRIN2B | -7.181081368 | Down |
| 6678 | SPARC | -5.871691208 | Down |
| 92745 | SLC38A5 | -5.409798061 | Down |
| 107984158 | LOC107984158 | -5.369470151 | Down |
| 7447 | VSNL1 | -5.31342666 | Down |
| 9358 | ITGBL1 | -5.105865851 | Down |
| 60489 | APOBEC3G | -5.027707987 | Down |
| 10561 | IFI44 | -4.797661758 | Down |
| 6515 | SLC2A3 | -4.500514925 | Down |
| Top 10 up-regulated DEGs |  |  |  |
| 1800 | DPEP1 | 8.227595159 | up |
| 11199 | ANXA10 | 6.660789481 | up |
| 7273 | TTN | 5.324255484 | up |
| 105372842 | LOC105372842 | 4.993254219 | up |
| 283383 | ADGRD1 | 4.035544471 | up |
| 54742 | LY6K | 3.783942219 | up |
| 945 | CD33 | 3.738284193 | up |
| 547 | KIF1A | 3.730261687 | up |
| 26240 | FAM50B | 3.545457095 | up |
| 57822 | GRHL3 | 3.477021065 | up |

### Enrichment analysis of DEGs in the ZKSCAN3-deficient colorectal cancer cell line

To determine the mechanism of DEGs by the effects of ZKSCAN3-deficient in colorectal cancer cell line, we performed the KEGG and GO analyses (Figure 2). We identified the top ten KEGG items which include “Herpes simplex virus 1 infection”, “Human papillomavirus infection”, “Epstein-Barr virus infection”, “Alcoholism”, “Viral carcinogenesis”, “Axon guidance”, “Antigen processing and presentation”, “ECM-receptor interaction”, “Pancreatic cancer”, and “Proteasome”. We also identified the top ten biological processes including “response to virus”, “kidney development”, “defense response to virus”, “defense response to symbiont”, “embryonic skeletal system morphogenesis”, “negative regulation of viral process”, “negative regulation of viral genome replication”, “regulation of viral genome replication”, “response to interferon-beta”, and “response to interferon-alpha”. We identified the top ten cellular components of GO including “cell-cell junction”, “transport vesicle”, “cell-substrate junction”, “collagen-containing extracellular matrix”, “endocytic vesicle”, “exocytic vesicle”, “contractile fiber”, “adherens junction”, “basement membrane”, and “platelet alpha granule lumen”. We then identified the top ten biological processes of GO, which contains “DNA-binding transcription activator activity”, “DNA-binding transcription activator activity, RNA polymerase II-specific”, “integrin binding”, “MHC protein complex binding”, “extracellular matrix structural constituent conferring tensile strength”, “oxidoreductase activity, acting on the aldehyde or oxo group of donors”, “calmodulin-dependent protein kinase activity”, “MHC class II protein complex binding”, “ion channel inhibitor activity”, and “type I transforming growth factor beta receptor binding”.

**Figure 2.**
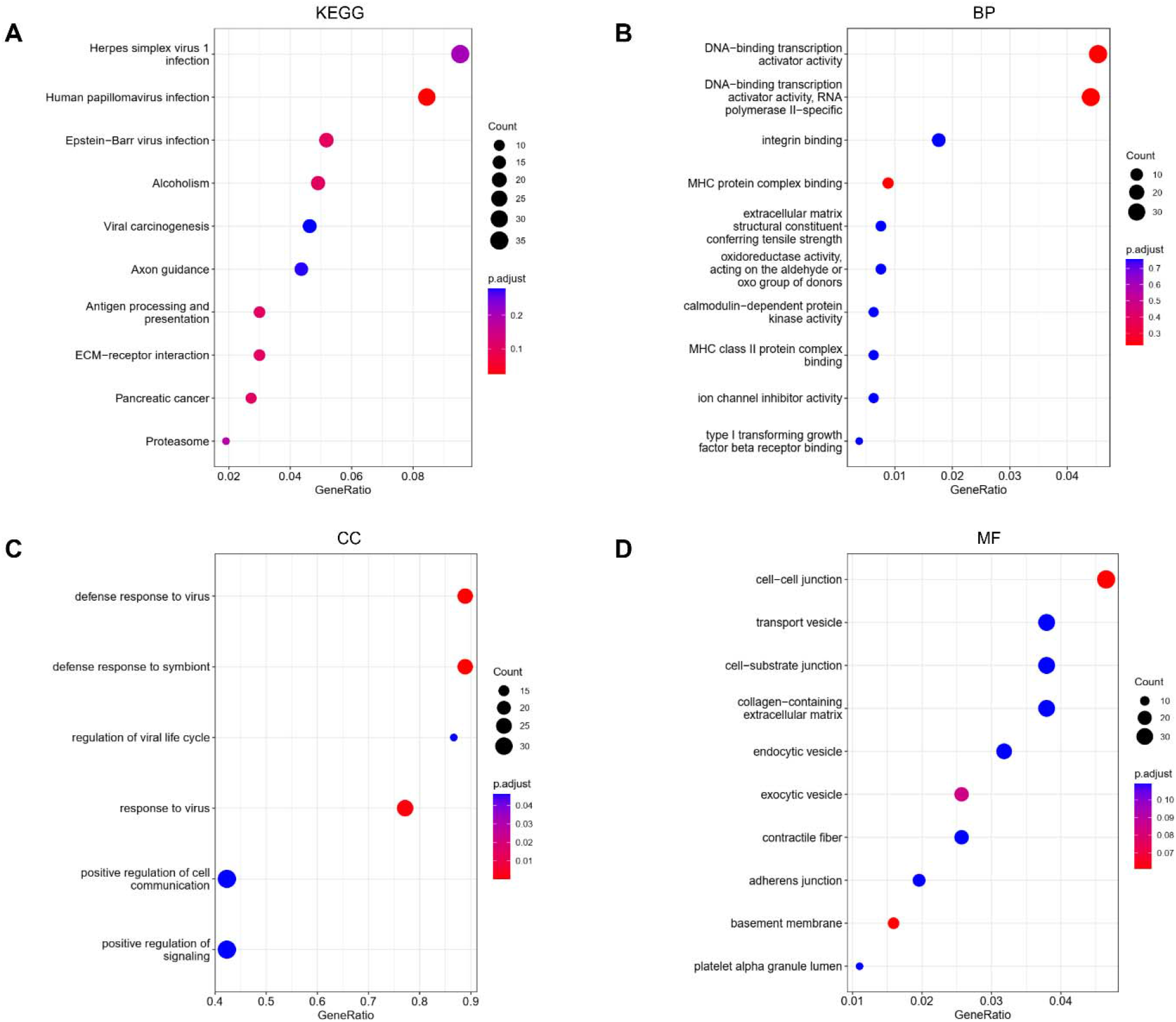
KEGG and GO analyses of DEGs between the WT and ZKSCAN3-deficient colorectal cancer cells. (A) KEGG analysis, (B) Biological processes, (C) Cellular components, (D) Molecular functions.

### Construction of PPI network in the ZKSCAN3-deficient colorectal cancer cell line

To understand the potential associations of DEGs, we constructed the PPI network by the Cytoscope software. The combined score > 0.2 was used as a cutoff to creat the PPI network by linking the 108 nodes and 93 edges. The Table 2 illustrated the top ten genes with the highest degree scores. The top two cluster modules were showed in Figure 3. We further evaluated the DEG and PPI by Reactome map (Figure 4) and figured out the top ten significant biological processes including “Activation of anterior HOX genes in hindbrain development during early embryogenesis”, “Activation of HOX genes during differentiation”, “Interferon alpha/beta signaling”, “RUNX2 regulates genes involved in cell migration”, “Interferon Signaling”, “Transcriptional regulation by RUNX2”, “Regulation of RUNX2 expression and activity”, “Unblocking of NMDA receptors, glutamate binding and activation”, “Ras activation upon Ca2+ influx through NMDA receptor”, and “Negative regulation of NMDA receptor-mediated neuronal transmission” (Supplemental Table S1).

**Figure 3.**
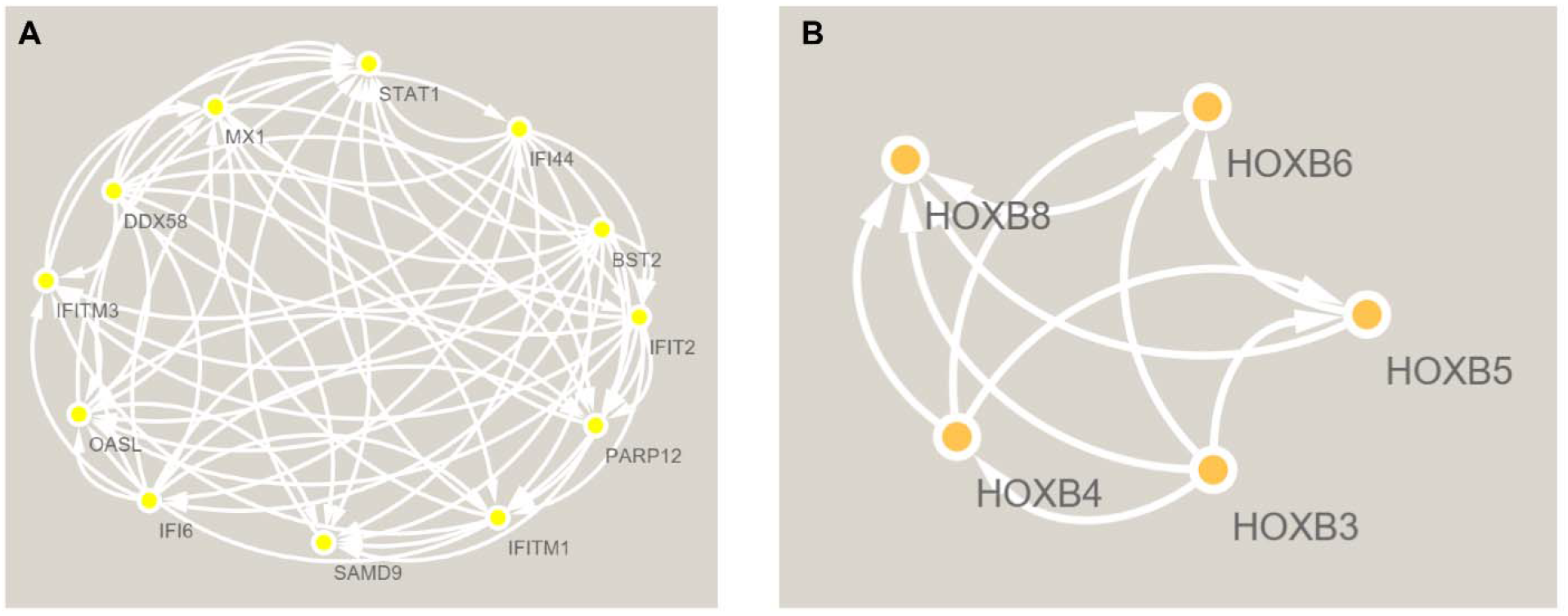
The PPI network analyses of DEGs between the WT and ZKSCAN3-deficient colorectal cancer cells. The cluster (A) and cluster (B) were constructed by MCODE.

**Figure 4.**
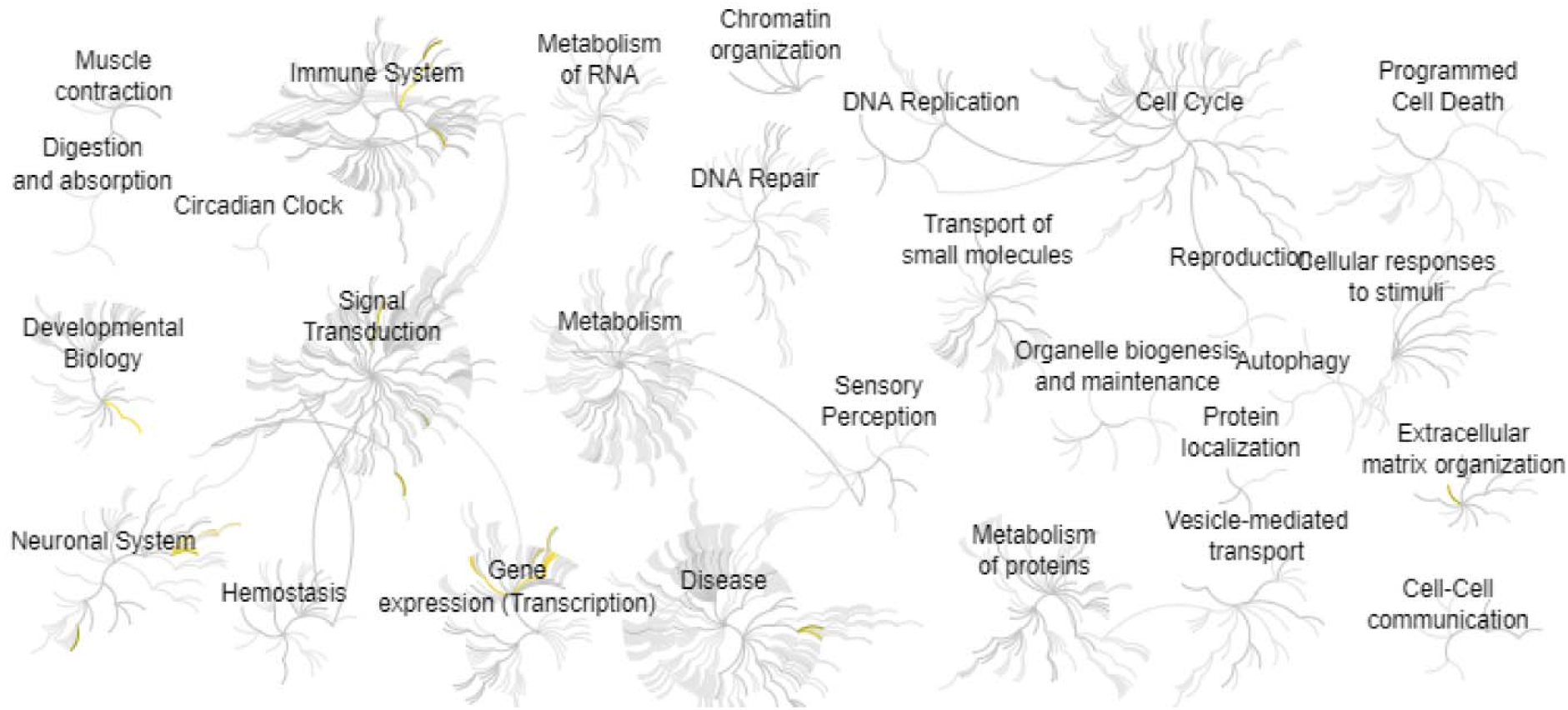
Reactome map representation of the significant biological processes between the WT and ZKSCAN3-deficient colorectal cancer cells.

**Table 2.** Top ten genes demonstrated by connectivity degree in the PPI network.

| **Gene symbol** | **Gene title** | **Degree** |
| --- | --- | --- |
| STAT1 | Signal Transducer And Activator Of Transcription 1 | 23 |
| MX1 | MX Dynamin Like Gtpase 1 | 17 |
| DDX58 | Dexd/H-Box Helicase 58 | 16 |
| PPARG | Peroxisome Proliferator Activated Receptor Gamma | 15 |
| EGFR | Epidermal Growth Factor Receptor | 15 |
| APP | Amyloid Beta Precursor Protein | 14 |
| BST2 | Bone Marrow Stromal Cell Antigen 2 | 14 |
| DLG4 | Discs Large MAGUK Scaffold Protein 4 | 14 |
| OASL | 2'-5'-Oligoadenylate Synthetase Like | 13 |
| IFIT2 | Interferon Induced Protein With Tetratricopeptide Repeats 2 | 12 |

## Discussion

Though there are two pathways (gatekeeper and caretaker) related to the progression of colorectal cancer^15^, the exact mechanism is still not clear. Thus, the treatment of colorectal cancer focusing on specific molecules attracts the most attention.

We found the immune defense response to virus and transcription activity are the mainly impacted processes in ZKSCAN3-deficient colorectal cancer cells. Lili Deng et al found the virus of Guang9 strain harboring IL-24 (VG9-IL-24) can repress the progression of colorectal cancer tumor by enhancing oncolysis and apoptosis as well as inducing the antitumor immune effects^16^. Adrian Jarzyński et al found the HPV and BKV infections can lead to the neoplastic process^17^. Shuaishuai Teng et al identified that the transcription factors FOXA2 and HNF1A can bind to the gained enhancers and activate the gene transcription to further promote the liver metastasis^18^. Moreover, ZKSCAN3 is a critical transcriptional repressor, which is higher expression in invasive compared with noninvasive colorectal cancer^19^.

Based on the PPI network, we identified a variety of interaction molecules in ZKSCAN3-deficient colorectal cancer cells. Chao Liu et al found that STAT1 regulates the inhibition of FOXM1 to enhance the gemcitabine sensitivity in pancreatic cancer^20^. Abrar I Aljohani et al found MX1 expression was related to the aggressiveness of cancer, including tumor size, grade, index scores, and high Ki67 expression, which is considered as an independent predictor of poor outcome in invasive breast cancer^21^. Yu-Chan Chang et al found the glucose transporter 4 promotes the oral cancer through the DDX58 signaling^22^. Andre Luiz Prezotto Villa et al found the expression of PPARG in colorectal cancer is related to the staging and clinical evolution^23^. Circadian gene clocks and its downstream molecules are related to many cell functions including metabolism, immune, proliferation, and differentiation^24–33^. PPARG is found to be a novel circadian transcription factor in adipogenesis and osteogenesis^34^. EGFR is related to the pathogenesis and progression of various cancers such as lung cancer^35^. GPCR and RGS proteins and downstream signaling pathways involve in the regulation of cell functions including proliferation, differentiation, migration, and secretion^36–40^. RGS proteins also play essential roles in a variety of diseases such as arthritis, osteoporosis, pain, and heart diseases^41–48^. RGS16 was found to be co-immunoprecipitated with EGFR and phosphorylated on both tyrosine residues after EGF stimulation in cancer cells^49^. APP was found to be increased in various cancer types, including lung and breast cancer^50^. BST2 is expressed in cancer cells, which enhances the progression of tumor and metastasis^51^. Keisuke Handa et al found DLG4 is a tumor inhibitor in the development of HPV-associated cervical cancers^52^. OASL proteins can mediate the cellular survival and affect the progression of cancer^53^. The downregulation of IFIT2 is related to the increased cell proliferation and metastasis in gastric cancer patients^54^.

In summary, our study identified the significant effects of ZKSCAN3 in colorectal cancer cells. The immune defense response to virus and transcription activity are the mainly impacted processes involved in the ZKSCAN3 mediated cancer cells. Therefore, this study may be beneficial to the treatment of colorectal cancer.

## Author Contributions

Zhiwen Qian, Tingxiang Chang, Tingting Zhang, Jing Wang: Methodology. Hanming Gu: Conceptualization, Writing and Editing.

## Funding

This work was not supported by any funding.

## Declarations of interest

There is no conflict of interest to declare.

## Notes

### Competing Interest Statement

The authors have declared no competing interest.

## References

[1] Haggar FA, Boushey RP: Colorectal cancer epidemiology: incidence, mortality, survival, and risk factors. Clin Colon Rectal Surg 2009, 22:191–7.

[2] Rawla P, Sunkara T, Barsouk A: Epidemiology of colorectal cancer: incidence, mortality, survival, and risk factors. Prz Gastroenterol 2019, 14:89–103.

[3] Sung H, Ferlay J, Siegel RL, Laversanne M, Soerjomataram I, Jemal A, Bray F: Global Cancer Statistics 2020: GLOBOCAN Estimates of Incidence and Mortality Worldwide for 36 Cancers in 185 Countries. CA Cancer J Clin 2021, 71:209–49.

[4] Johnson CM, Wei C, Ensor JE, Smolenski DJ, Amos CI, Levin B, Berry DA: Meta-analyses of colorectal cancer risk factors. Cancer Causes Control 2013, 24:1207–22.

[5] Fares J, Fares MY, Khachfe HH, Salhab HA, Fares Y: Molecular principles of metastasis: a hallmark of cancer revisited. Signal Transduct Target Ther 2020, 5:28.

[6] Jasperson KW, Tuohy TM, Neklason DW, Burt RW: Hereditary and familial colon cancer. Gastroenterology 2010, 138:2044–58.

[7] Huang M, Chen Y, Han D, Lei Z, Chu X: Role of the zinc finger and SCAN domain-containing transcription factors in cancer. Am J Cancer Res 2019, 9:816–36.

[8] Ola MS, Nawaz M, Ahsan H: Role of Bcl-2 family proteins and caspases in the regulation of apoptosis. Mol Cell Biochem 2011, 351:41–58.

[9] Yang L, Zhang L, Wu Q, Boyd DD: Unbiased screening for transcriptional targets of ZKSCAN3 identifies integrin beta 4 and vascular endothelial growth factor as downstream targets. J Biol Chem 2008, 283:35295–304.

[10] Yu G, Wang LG, Han Y, He QY: clusterProfiler: an R package for comparing biological themes among gene clusters. OMICS 2012, 16:284–7.

[11] Hanming G: nuotrophils arthritis. Research Square 2021.

[12] Jing L, Letian W, Hanming G: Identification of driver genes and biological signaling for alcoholic myopathy. Research Square 2021.

[13] Li J, Wang W, Gu H: Identification of biological processes and signaling pathways for the knockout of REV-ERB in mouse brain. bioRxiv 2021:2021.11.22.469579.

[14] Yuan G: Identification of biomarkers and pathways of mitochondria in sepsis patients. bioRxiv 2021:2021.03.29.437586.

[15] Russo A, Zanna I, Tubiolo C, Migliavacca M, Bazan V, Latteri MA, Tomasino RM, Gebbia N: Hereditary common cancers: molecular and clinical genetics. Anticancer Res 2000, 20:4841–51.

[16] Deng L, Yang X, Fan J, Ding Y, Peng Y, Xu D, Huang B, Hu Z: IL-24-Armed Oncolytic Vaccinia Virus Exerts Potent Antitumor Effects via Multiple Pathways in Colorectal Cancer. Oncol Res 2021, 28:579–90.

[17] Jarzynski A, Zajac P, Zebrowski R, Boguszewska A, Polz-Dacewicz M: Occurrence of BK Virus and Human Papilloma Virus in colorectal cancer. Ann Agric Environ Med 2017, 24:440–5.

[18] Teng S, Li YE, Yang M, Qi R, Huang Y, Wang Q, Zhang Y, Chen S, Li S, Lin K, Cao Y, Ji Q, Gu Q, Cheng Y, Chang Z, Guo W, Wang P, Garcia-Bassets I, Lu ZJ, Wang D: Tissue-specific transcription reprogramming promotes liver metastasis of colorectal cancer. Cell Res 2020, 30:34–49.

[19] Yang L, Hamilton SR, Sood A, Kuwai T, Ellis L, Sanguino A, Lopez-Berestein G, Boyd DD: The previously undescribed ZKSCAN3 (ZNF306) is a novel “driver” of colorectal cancer progression. Cancer Res 2008, 68:4321–30.

[20] Liu C, Shi ì, Li Q, Li Z, Lou C, Zhao Q, Zhu Y, Zhan F, Lian J, Wang B, Guan X, Fang L, Li Z, Wang Y, Zhou B, Yao Y, Zhang Y: STAT1-mediated inhibition of FOXM1 enhances gemcitabine sensitivity in pancreatic cancer. Clin Sci (Lond) 2019, 133:645–63.

[21] Aljohani Ai, Joseph C, Kurozumi S, Mohammed OJ, Miligy IM, Green AR, Rakha EA: Myxovirus resistance 1 (MX1) is an independent predictor of poor outcome in invasive breast cancer. Breast Cancer Res Treat 2020, 181:541–51.

[22] Chang YC, Chi LH, Chang WM, Su CY, Lin YF, Chen CL, Chen MH, Chang PM, Wu AT, Hsiao M: Glucose transporter 4 promotes head and neck squamous cell carcinoma metastasis through the TRIM24-DDX58 axis. J Hematol Oncol 2017, 10:11.

[23] Villa ALP, Parra RS, Feitosa MR, Camargo HP, Machado VF, Tirapelli D, Rocha J, Feres O: PPARG expression in colorectal cancer and its association with staging and clinical evolution. Acta Cir Bras 2020, 35:e202000708.

[24] Yuan G, Hua B, Cai T, Xu L, Li E, Huang Y, Sun N, Yan Z, Lu C, Qian R: Clock mediates liver senescence by controlling ER stress. Aging 2017, 9:2647–65.

[25] Yuan G, Xu L, Cai T, Hua B, Sun N, Yan Z, Lu C, Qian R: Clock mutant promotes osteoarthritis by inhibiting the acetylation of NFkappaB. Osteoarthritis Cartilage 2019, 27:922–31.

[26] Zhu Z, Xu L, Cai T, Yuan G, Sun N, Lu C, Qian R: Clock represses preadipocytes adipogenesis via GILZ. J Cell Physiol 2018, 233:6028–40.

[27] Fan XF, Wang XR, Yuan GS, Wu DH, Hu LG, Xue F, Gong YS: [Effect of safflower injection on endoplasmic reticulum stress-induced apoptosts in rats with hypoxic pulmonary hypertension], Zhongguo Ying Yong Sheng Li Xue Za Zhi 2012, 28:561–7.

[28] Yuan G, Hua B, Yang Y, Xu L, Cai T, Sun N, Yan Z, Lu C, Qian R: The Circadian Gene Clock Regulates Bone Formation Via PDIA3. J Bone Miner Res 2017, 32:861–71.

[29] Xu L, Cheng Q, Hua B, Cai T, Lin J, Yuan G, Yan Z, Li X, Sun N, Lu C, Qian R: Circadian gene Clock regulates mitochondrial morphology and functions by posttranscriptional way. bioRxiv 2018:365452.

[30] Zhu Z, Hua B, Xu L, Yuan G, Li E, Li X, Sun N, Yan Z, Lu C, Qian R: CLOCK promotes 3T3-L1 cell proliferation via Wnt signaling. IUBMB Life 2016, 68:557–68.

[31] Cai T, Hua B, Luo D, Xu L, Cheng Q, Yuan G, Yan Z, Sun N, Hua L, Lu C: The circadian protein CLOCK regulates cell metabolism via the mitochondrial carrier SLC25A10. Biochim Biophys Acta Mol Cell Res 2019, 1866:1310–21.

[32] Zhu Z, Hua B, Shang Z, Yuan G, Xu L, Li E, Li X, Sun N, Yan Z, Qian R, Lu C: Altered Clock and Lipid Metabolism-Related Genes in Atherosclerotic Mice Kept with Abnormal Lighting Condition. Biomed Res Int 2016, 2016:5438589.

[33] Mao SZ, Fan XF, Xue F, Chen R, Chen XY, Yuan GS, Hu LG, Liu SF, Gong YS: Intermedin modulates hypoxic pulmonary vascular remodeling by inhibiting pulmonary artery smooth muscle cell proliferation. Pulm Pharmacol Ther 2014, 27:1–9.

[34] Kawai M, Rosen CJ: PPARgamma: a circadian transcription factor in adipogenesis and osteogenesis. Nat Rev Endocrinol 2010, 6:629–36.

[35] da Cunha Santos G, Shepherd FA, Tsao MS: EGFR mutations and lung cancer. Annu Rev Pathol 2011, 6:49–69.

[36] Doze VA, Perez DM: GPCRs in stem cell function. Prog Mol Biol Transl Sci 2013, 115:175–216.

[37] Yuan G, Fu C, Yang ST, Yuh DY, Hajishengallis G, Yang S: RGS12 Drives Macrophage Activation and Osteoclastogenesis in Periodontitis. J Dent Res 2021:220345211045303.

[38] Yuan G, Yang S, Yang S: Macrophage RGS12 contributes to osteoarthritis pathogenesis through enhancing the ubiquitination. Genes & Diseases 2021.

[39] Fu C, Yuan G, Yang ST, Zhang D, Yang S: RGS12 Represses Oral Cancer via the Phosphorylation and SUMOylation of PTEN. J Dent Res 2020:22034520972095.

[40] Zhang X, Eggert US: Non-traditional roles of G protein-coupled receptors in basic cell biology. Mol Biosyst 2013, 9:586–95.

[41] Yuan G, Huang Y, Yang S-t, Ng A, Yang S: RGS12 inhibits the progression and metastasis of multiple myeloma by driving M1 macrophage polarization and activation in the bone marrow microenvironment. Cancer Commun, n/a.

[42] Yuan G, Yang S, Ng A, Fu C, Oursler MJ, Xing L, Yang S: RGS12 Is a Novel Critical NF-kappaB Activator in Inflammatory Arthritis. iScience 2020, 23:101172.

[43] Tsang S, Woo AY, Zhu W, Xiao RP: Deregulation of RGS2 in cardiovascular diseases. Front Biosci (Schol Ed) 2010, 2:547–57.

[44] Tsingotjidou A, Nervina JM, Pham L, Bezouglaia O, Tetradis S: Parathyroid hormone induces RGS-2 expression by a cyclic adenosine 3’,5’-monophosphate-mediated pathway in primary neonatal murine osteoblasts. Bone 2002, 30:677–84.

[45] Yuan G, Yang S, Gautam M, Luo W, Yang S: Macrophage regulator of G-protein signaling 12 contributes to inflammatory pain hypersensitivity. Ann Transl Med 2021, 9:448.

[46] Avrampou K, Pryce KD, Ramakrishnan A, Sakloth F, Gaspari S, Serafini RA, Mitsi V, Polizu C, Swartz C, Ligas B, Richards A, Shen L, Carr FB, Zachariou V: RGS4 Maintains Chronic Pain Symptoms in Rodent Models. J Neurosci 2019, 39:8291–304.

[47] Yuan G, Yang S, Yang S, Ng A, Oursler MJ: RGS12 is a critical proinflammatory factor in the pathogenesis of inflammatory arthritis via acting in Cox2-RGS12-NF kappa B pathway activation loop. J Bone Miner Res: WILEY 111 RIVER ST, HOBOKEN 07030-5774, NJ USA, 2019. pp. 147–.

[48] Yuan G, Yang S, Liu M, Yang S: RGS12 is required for the maintenance of mitochondrial function during skeletal development. Cell Discov 2020, 6:59.

[49] Derrien A, Druey KM: RGS16 function is regulated by epidermal growth factor receptor-mediated tyrosine phosphorylation. J Biol Chem 2001, 276:48532–8.

[50] Wu X, Chen S, Lu C: Amyloid precursor protein promotes the migration and invasion of breast cancer cells by regulating the MAPK signaling pathway. Int J Mol Med 2020, 45:162–74.

[51] Mahauad-Fernandez WD, DeMali KA, Olivier AK, Okeoma CM: Bone marrow stromal antigen 2 expressed in cancer cells promotes mammary tumor growth and metastasis. Breast Cancer Res 2014, 16:493.

[52] Handa K, Yugawa T, Narisawa-Saito M, Ohno S, Fujita M, Kiyono T: E6AP-dependent degradation of DLG4/PSD95 by high-risk human papillomavirus type 18 E6 protein. J Virol 2007, 81:1379–89.

[53] Choi UY, Kang JS, Hwang YS, Kim YJ: Oligoadenylate synthase-like (OASL) proteins: dual functions and associations with diseases. Exp Mol Med 2015, 47:e144.

[54] Pidugu VK, Pidugu HB, Wu MM, Liu CJ, Lee TC: Emerging Functions of Human IFIT Proteins in Cancer. Front Mol Biosci 2019, 6:148.

